# Degrees of compositional shift in tree communities vary along a gradient of temperature change rates over one decade: Application of an individual-based temporal beta-diversity concept

**DOI:** 10.1101/2020.01.30.920660

**Authors:** Ryosuke Nakadai

## Abstract

Temporal patterns in communities have gained widespread attention recently, to the extent that temporal changes in community composition are now termed “temporal beta-diversity”. Previous studies of beta-diversity have made use of two classes of dissimilarity indices: incidence-based (e.g., Sørensen and Jaccard dissimilarity) and abundance-based (e.g., Bray–Curtis and Ružička dissimilarity). However, in the context of temporal beta-diversity, the persistence of identical individuals and turnover among other individuals within the same species over time have not been considered, despite the fact that both will affect compositional changes in communities. To address this issue, I propose new index concepts for beta-diversity and the relative speed of compositional shifts in relation to individual turnover based on individual identity information. Individual-based beta-diversity indices are novel dissimilarity indices that consider individual identity information to quantitatively evaluate temporal change in individual turnover and community composition. I applied these new indices to individually tracked tree monitoring data in deciduous and evergreen broad-leaved forests across the Japanese archipelago with the objective of quantifying the effect of climate change trends (i.e., rates of change of both annual mean temperature and annual precipitation) on individual turnover and compositional shifts at each site. A new index explored the relative contributions of mortality and recruitment processes to temporal changes in community composition. Clear patterns emerged showing that an increase in the temperature change rate facilitated the relative contribution of mortality components. The relative speed of compositional shift increased with increasing temperature change rates in deciduous forests but decreased with increasing warming rates in evergreen forests. These new concepts provide a way to identify novel and high-resolution temporal patterns in communities.

## Introduction

All living organisms have finite life spans. Each individual tries to leave offspring before its death. The persistence of individuals and their reproduction in the past shape the current diversity patterns through temporal dynamics of populations, communities, and ecosystems. The effects of recent anthropogenically induced climate change on local and global biodiversity have received considerable attention (Parmesan and Yohe 2003, Chen et al. 2011). Specifically, intensive efforts have focused on biodiversity loss (i.e., alpha- and gamma-diversity) and compositional shifts (i.e., beta-diversity) within the context of biodiversity conservation (e.g., Dornelas et al. 2014, Valtonen et al. 2016, Hillebrand et al. 2017). Ideally, long-term biodiversity monitoring programs from multiple countries employing consistent methodologies should be used to identify the effects of global climate change. However, multi-decade monitoring programs remain very rare (Magurran et al. 2010; but see Valtonen et al. 2016).

Whittaker (1960, 1972) developed the concept of beta-diversity, which he defined as the variation in community composition among sites in a region. Since then, many new beta-diversity indices have been developed, and attempts have been made to relate the properties of beta-diversity and dissimilarity indices to background ecological processes (Harrison et al. 1992, Williams 1996, Lennon et al. 2001, Tuomisto and Ruokolainen 2006, Baselga 2010, Legendre 2010; reviewed by Koleff et al. 2003, Anderson et al. 2011, Jost et al. 2011, Legendre and De Cáceres 2013). More attention has been focused recently on temporal changes in community composition in either single sites or in a series of sites surveyed repeatedly over time (Magurran 2011, Bahram et al. 2015, Matsuoka et al. 2016, Legendre and Condit 2019). Temporal changes in community composition are now termed “temporal beta-diversity” (Hatosy et al. 2013, Legendre and Gauthier 2014, Shimadzu et al. 2015). Legendre (2019) elegantly partitioned temporal beta-diversity into loss and gain components of the compositional changes in each community, thereby facilitating mechanistic understanding of the temporal changes in community composition. Furthermore, recent studies have started to show that the temporal beta-diversity patterns differ among sites (Legendre and Condit 2019, Magurran et al. 2019). Environmental changes, including anthropogenically induced shifts such as global climate change, are likely factors that drive spatial differences in temporal beta-diversity (Legendre 2019).

The persistence of individuals over time is an important component of the temporal beta-diversity concept that has not been explicitly considered previously. The persistence of individuals contributes to a static equilibrium in community composition. By contrast, turnover of different individuals belonging to the same species, even when the community composition is stable over time, contributes to a dynamic equilibrium in community composition. Therefore, conventional temporal beta-diversity is zero when the community composition is at equilibrium. However, it can be difficult to detect differences in turnover within the same species and the speed or frequency of compositional changes in communities differs between sites with high and low individual turnover rates across time. Therefore, an appropriate evaluation of temporal changes in community composition could benefit from considering the fates of identifiable individuals in a community over time. Here, I propose new indices of temporal beta-diversity that take into account individual identity information. These new coefficients are individual-based in contrast to others that are incidence-based (e.g., the Sørensen index; Sørensen 1948), or abundance-based (e.g., the Bray–Curtis index; Bray and Curtis 1957). My new indices quantitatively evaluate the temporal changes in community composition by considering individual identity information and reveal the relative contribution of different background processes to these temporal changes. I also propose a new index to quantify the relative speed of change away from compositional equilibrium in a community relative to the speed of individual turnover. First, I calculated these indices for 16 forest inventory datasets compiled across the Japanese archipelago. These datasets have individual identity information for two time periods. I then aimed to empirically quantify the extent of the compositional shift response to climate change by excluding the effect of differences in the speed of individual turnover among site. The indices identified the effects of climate change on the composition of tree communities on a decadal time scale.

## Materials and Methods

### New indices of temporal beta-diversity and new statistics to evaluate the relative speed of compositional shift in relation to individual turnover

Two types of dissimilarity indices have been proposed in previous studies: incidence-based and abundance-based (Baselga et al. 2013). Specifically, the widely used Bray– Curtis dissimilarity index (Bray and Curtis 1957, which was proposed earlier as a proportional difference index by Odum 1950), is an abundance-based extension of the Sørensen index (Legendre and Legendre 1998, Baselga et al. 2013). I have excluded details of incidence-based indices that are tangential to the main topic in my study (see Anderson et al. 2011, Jost et al. 2011, Baselga et al. 2013). The conceptual basis for conventional beta-diversity indices and my new procedures relies on the fact that the intersection (*A* component) and the relative complements (*B* and *C* components) for species abundance can be formulated as follows: 

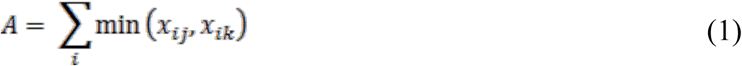

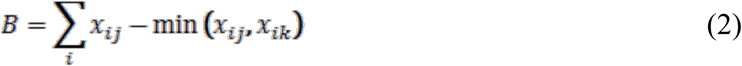

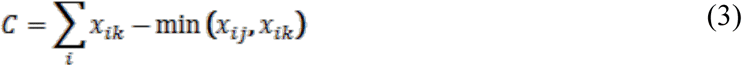

where *x*_*ij*_ is the abundance of species *i* at site *j*, and *x*_*ik*_ is the abundance of species *i* at site *k*. Therefore, *A* is the number of individuals of each species that occur at both sites *j* and *k*, while *B* and *C* are the numbers of individuals that are unique to sites *j* and *k*, respectively. This formulation of the Bray–Curtis (*d*_*BC*_) index is expressed as follows: 

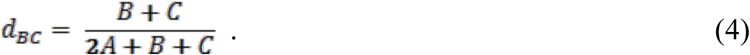

In the temporal beta-diversity concept, the differences between data vectors correspond to observations made at time 1 (abbreviated T1) and time 2 (T2), which are referred to sites *k* and *j* in the spatial beta-diversity concept. The *A, B*, and *C* components are described as follows: 

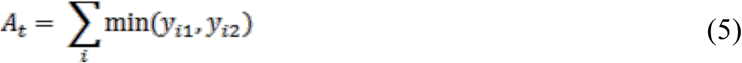

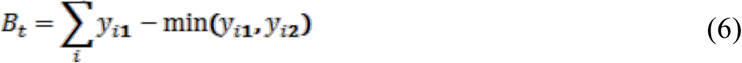

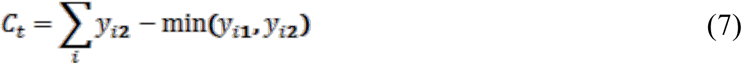

where *y*_*i1*_ is the abundance of species *i* at time T1 at a site, and *y*_*i2*_ is the abundance of species *i* at time T2 at the site. Therefore, *A*_*t*_ is the number of individuals of each species that occur at both times T1 and T2, whereas *B*_*t*_ and *C*_*t*_ are the numbers of individuals that are unique to times T1 and T2, respectively. The temporal beta-diversity of the Bray–Curtis (*d*_*t*.*BC*_) index is expressed as follows: 

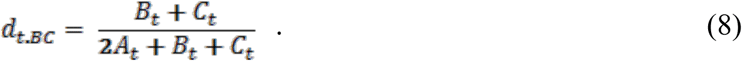

Using Equation 8, Legendre (2019) partitioned temporal beta-diversity indices into loss and gain components because a change through time is directional; species abundances would identify losses and/or gains between T1 and T2 as follows: 

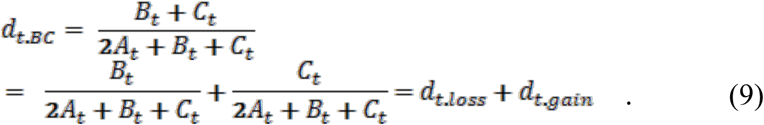

Legendre and his collaborators applied the loss/gain partitioning procedure for temporal beta-diversity to many empirical datasets (e.g., Kuczynski et al. 2018, Brice et al. 2019), and constructed a new analytical trend in temporal beta-diversity.

### Individual-based components and novel indices

In this study, I developed the concept of individual-based temporal beta-diversity based on Bray–Curtis dissimilarity through extension of the abundance-based temporal beta-diversity indices. Traditionally, the equilibrium and non-equilibrium elements in community ecology indicate properties of the whole community that are directly related to species coexistence (Pickett 1980). However, some individuals replace others, and thus ecological communities are always dynamic and vary by at least some degree in space and time (Mori et al. 2018, Ryo et al. 2019). In addition, some individuals of long-lived species (e.g., trees and vertebrates) persist over the long term, whereas individuals of short-lived species are replaced by conspecifics or different species. Therefore, it should be possible to categorize the components of community dynamics into persistence (*P*), and those that change over time, i.e., mortality and recruitment (*M* and *R*, respectively). M refers to individuals occurring at T1 that die before T2, thereby representing the death of individuals during the time period between T1 and T2 (Fig. 1). *R* identifies individuals that were not alive at T1 but were subsequently found at T2, thereby representing recruitment during the period between T1 and T2. Component *P* refers to the persistence of individuals from T1 to T2 (Fig. 1). Although *P, M*, and *R* relate to components *A, B*, and *C*, respectively, here I have used different characters to emphasize the processes represented by each component. The respective abundances were calculated as follows:

**Figure 1.**
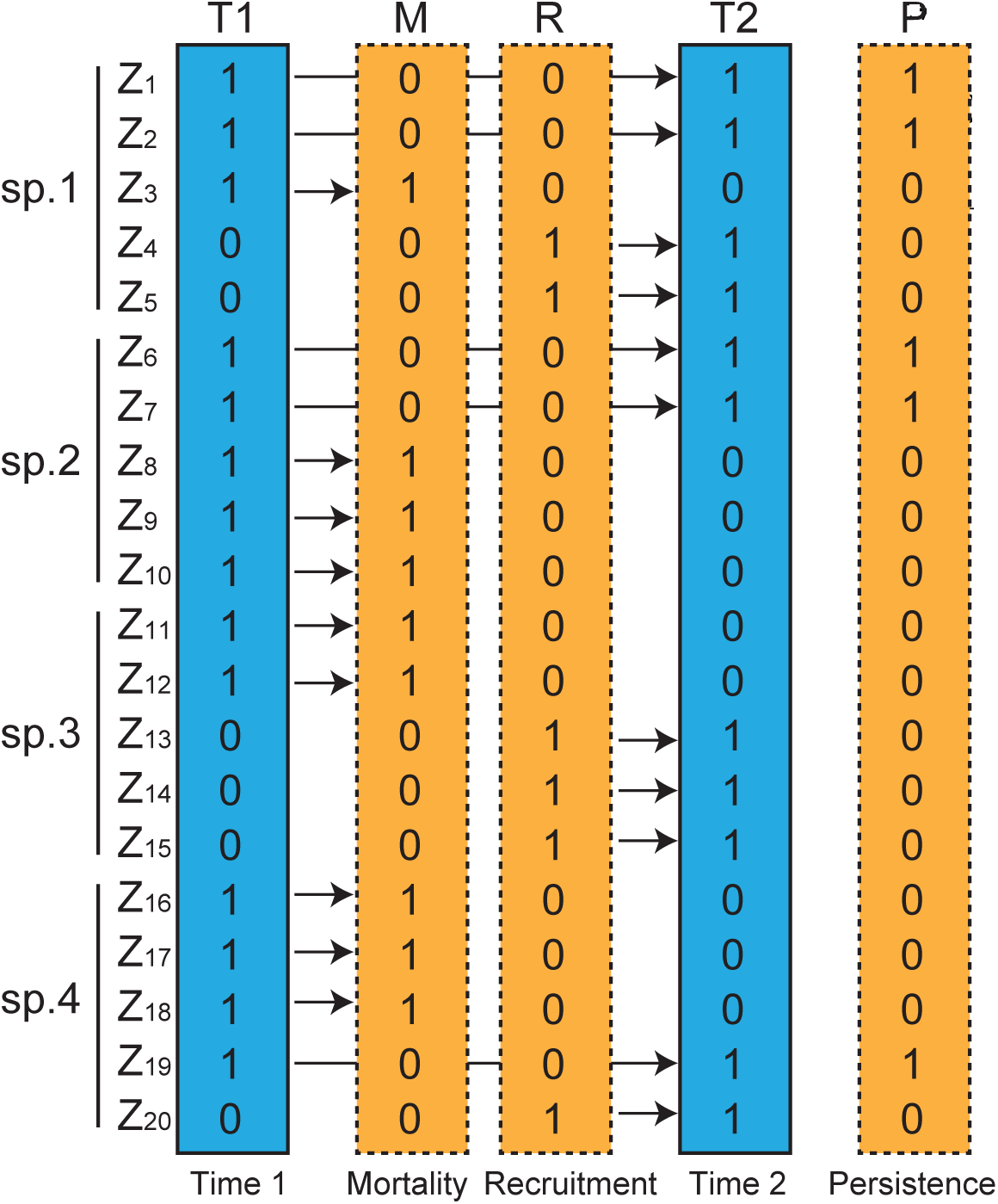
Schematic representation of individual-based temporal beta-diversity. T1 and T2 refer to the communities at times 1 (abbreviated T1) and 2 (T2), respectively. Both communities comprise four species (sp. 1, 2, 3, and 4) and all individuals are identified separately (*Zl*). M refers to individuals in existence at T1 but dead before T2, thereby separately identifying the deaths of individuals during the period between T1 and T2. R identifies individuals that were not alive at T1, but found at T2, thereby identifying the recruitment of individuals during the period between T1 and T2. P refers to individuals that persisted from T1 to T2, thus both T1 and T2 are 1. The arrows indicate the temporal trajectory for each individual; specifically, when they are present or absent in communities between T1 and T2.

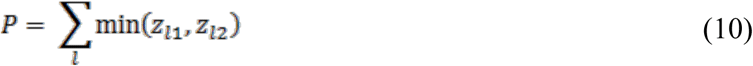

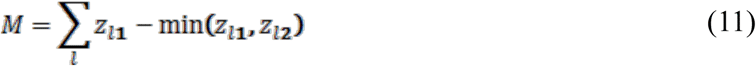

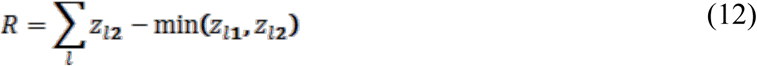

where *z*_*l1*_ is individual *l* at time T1, and *z*_*l2*_ is individual *l* at time T2. Both *z*_*l1*_ and *z*_*l2*_ take only *1* or 0, i.e., presence or absence. Therefore, *P* is the number of individuals present at both T1 and T2, whereas *M* and *R* are the numbers of individuals that are unique to T1 or T2, respectively. This formulation of the individual-based temporal beta-diversity index (*d*_*MR*_) can be expressed as follows: 

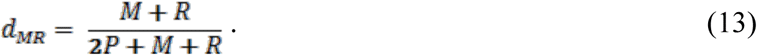

Furthermore, this individual-based temporal beta-diversity can be partitioned into the relative contributions of mortality (*d*_*M*_) and recruitment (*d*_*R*_) processes using a similar procedure for the loss and gain component-partitioning of the temporal beta-diversity in Equation 9 (Legendre 2019) as follows: 

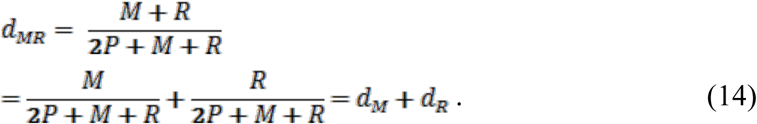

In some cases, there is turnover of different individuals belonging to the same species even when the community composition is stable over time, which contributes to a dynamic compositional equilibrium across time periods. Therefore, the components that change over time (*M* and *R*) are also separated into two components; compositional equilibrium (*E*) and shift (*B*_*t*_ and *C*_*t*_), respectively. Component *M* can be partitioned into lost individuals that contribute to compositional change in a community (*B*_t_) and lost individuals that are replaced by conspecifics, contributing substantially to equilibrium in community composition (*E*_*loss*_). Similarly, component *R* can be partitioned into gained individuals that contribute to compositional change in a community (*C*_*t*_), and gained individuals replacing conspecifics, thereby contributing substantially to equilibrium in community composition (*E*_*gain*_). By definition, the numbers of *E*_*loss*_ and *E*_*gain*_ individuals are identical; hence, I replaced both *E*_*loss*_ and *E*_*gain*_ with the coefficient *E*, the component that contributes to dynamic compositional equilibrium. The formulation is as follows: 

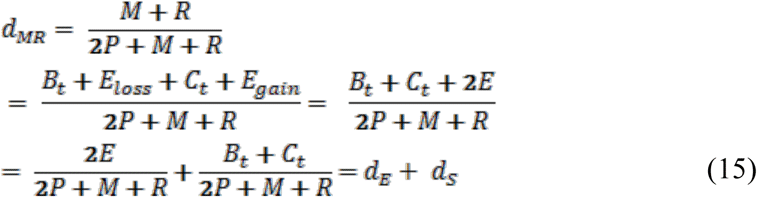

where *d*_*E*_ and *d*_*S*_ indicate the contributions of equilibrium and shift to individual-based temporal beta-diversity. I note that *E* is equal to the *A*_*t*_ component minus the *P* component.

Both *d*_*M*_/*d*_*R*_ and *d*_*E*_/*d*_*S*_ are indicators of the relative contribution to individual-based temporal beta-diversity (*d*_*MR*_); they represent rates of mortality vs. recruitment processes, and compositional equilibrium to shift, respectively. Furthermore, the components *M* and *R* can also be analyzed in the same manner as community data (Fig. 1). Therefore, the calculation of Bray–Curtis dissimilarity between communities using mortality and recruitment is as follows: 

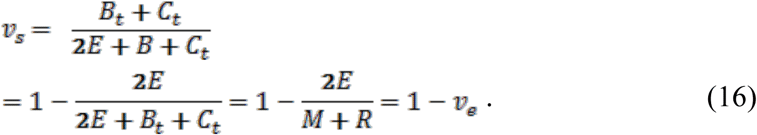

*v*_*s*_ indicates the speed of compositional shifts in a community relative to the total speed of individual turnover through mortality and recruitment processes. If *v*_*s*_ is 0, then individual turnover would not contribute to compositional shifts in a community, i.e., the community composition of mortality and recruitment are identical and an equilibrium state exists. If *v*_*s*_ is 1, all individual turnovers contribute to compositional shifts, and the community composition of mortality and recruitment are completely different. It is also possible to calculate *v*_*s*_ as *d*_*t.BC*_/*d*_*MR*_. Although I did not use *v*_*e*_ for analyses in the present study, *v*_*e*_ should be an index of dynamic compositional equilibrium. The relative contributions of compositional equilibrium and shift (*d*_*E*_/*d*_*S*_) are formulated as follows: 

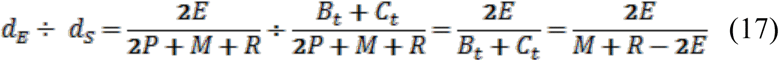

Because “2*E*” in the denominator is simply the difference between *v*_*e*_ (equation 16) and *d*_*E*_/*d*_*S*_ (equation 17), the two are strongly correlated (Table S1), but these two indices have different meanings.

In my main analyses, I focused only on an extension of Bray–Curtis dissimilarity. I also summarized explanations of individual-based beta-diversity using the Ružička dissimilarity index (Ružička 1958), which was identified by Podani et al. (2013) and Legendre (2014) as another appropriate index for beta-diversity studies. The Ružička dissimilarity index is also an abundance-based extension of the Jaccard dissimilarity index (Jaccard 1901). The explanations of individual-based beta-diversity using the Ružička dissimilarity index are described Supplementary text 1.

### Applying Japanese forest monitoring datasets to testing patterns in temporal beta-diversity indices along climate change gradients

I predicted the indices of individual-based beta-diversity and the relative speed of compositional shift variation along degrees of climate change rate per year, i.e., average speed of local climate change. I applied the new indices using individual-tracked monitoring data from forests in Japan. Two types of temporal change in tree communities may result from climate change. One is due to changes in the obligate speed of individual turnover and the balance of mortality and recruitment, e.g., climate warming facilitates the faster growth and shorter life span of each individual tree as a physiological process within individuals (Körner 2017, Munné-Bosch 2018). The other is due to the relative speed of compositional change, e.g., simple compositional shifts driven by climate change as an ecological process among individuals (i.e., competitive interactions and adaptation to the environment). Those can be referred to as *d*_*MR*_, including the components (*d*_*M*_, *d*_*R*_) and *v*_*s*_, respectively.

The Japanese archipelago comprises a long chain of continental islands lying off the eastern coast of Asia; it is recognized as a hotspot of biodiversity (Mittermeier et al. 2011). The mean temperatures of the coldest months range from –18.4 to 22.3°C, mean temperatures of the warmest months from 6.2 to 29.2°C, and the annual precipitation ranges from 700.4 to 4,477.2 mm (Japan Meteorological Agency, 1981–2010; http://www.jma.go.jp/jma/indexe.html). Typical forest biomes are classified into four types: alpine meadow/creeping pine zone, subalpine coniferous forest zone, deciduous broad-leaved forest zone, and evergreen broad-leaved forest zone. The empirical tree census dataset used in this study was obtained from the forest plots used in the Monitoring Sites 1000 Project; this project was launched by the Ministry of the Environment, Japan (see Ishihara et al. 2011 for details). The monitoring sites include five types of forest: deciduous broad-leaved, evergreen broad-leaved, mixed broad-leaved and coniferous, evergreen coniferous, and artificial forest (Ishihara et al., 2011, Suzuki et al. 2015). The full dataset is available on the website of the Biodiversity Center of Japan (http://www.biodic.go.jp/moni1000/index.html, in Japanese: accessed on November 11, 2019), and a subset of the data has been published as a data article (Ishihara et al. 2011).

To apply the individual-based temporal beta-diversity approach, I selected 16 forest plots. These included nine plots of deciduous broad-leaved forest and seven plots of evergreen broad-leaved forest (Fig. 2, Table S2) that met the following criteria: species level identifications had been performed in each plot, the plots had been surveyed one decade after a previous survey (in the period 2005–2008 to 2015–2018), individual identity information was recorded throughout the two time periods, plot size was 1.0 ha (100 × 100 m), and sufficient plot sample sizes in various environments within each forest type were available for use in analyses. In this analysis, I selected tree individuals with stem girths at breast height (GBH) ≥ 15.7 cm, which was approximately equivalent to a diameter at breast height (DBH) ≥ 5 cm; these were considered target individuals in the communities. I organized the data at the level of individual trees before the analyses, the original forest datasets had been compiled at the stem level, using taxonomic information provided in the Ylist (Yonekura and Kajita 2003). The data I analyzed included 18,749 individuals belonging to 254 species found in the 16 plots (Fig. 2).

**Figure 2.**
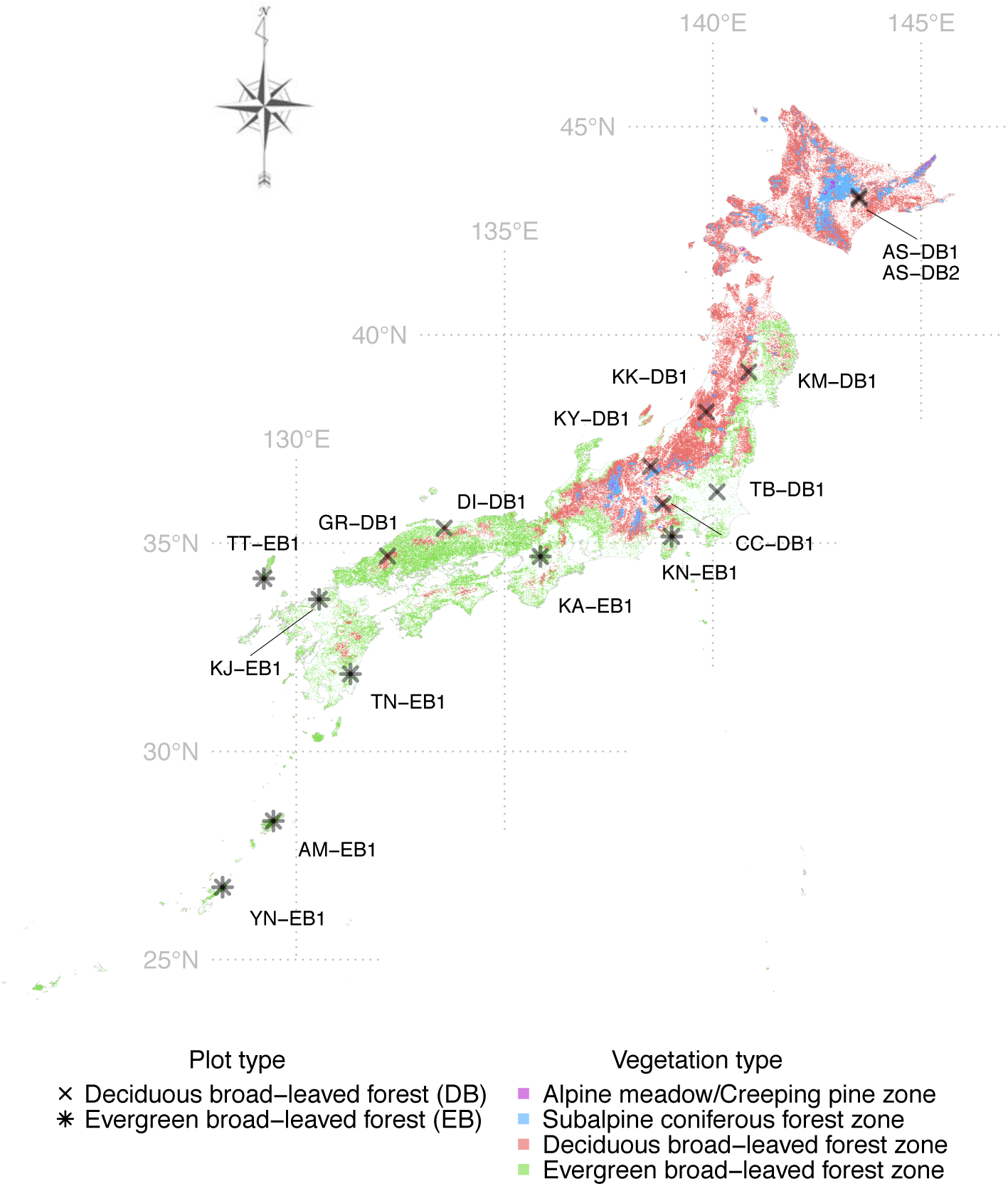
Locations of the study sites within the Japanese archipelago. The crosses and asterisks indicate the locations of targeted plots in deciduous broad-leaved forest (DB) and evergreen broad-leaved forest (EB) included in the present study. The lettering adjacent to each symbol is the plot identification code (ID). Detailed plot information is listed in Table S3. The colors indicate four major vegetation types: purple, Alpine meadow or creeping pine zone; blue, subalpine coniferous forest zone; red, deciduous broad-leaved forest zone; green, evergreen broad-leaved forest zone. The information on vegetation type at a 1.0-km grid scale was obtained from the results of the fifth censuses (1992–1996) of the National Survey of the Natural Environment in Japan (http://www.biodic.go.jp/dload/mesh_vg.html; in Japanese, accessed December 12, 2019).

### Climate variables

To obtain meteorological observation data and their changes over a decade, I searched for the meteorological station closest to each of the forest plots in the Japanese Automated Meteorological Data Acquisition System database (Japan Meteorological Agency; see details in Table S2, Fig. S1). Specifically, I extracted the monthly mean and total precipitation data (accessed December 12, 2019), and calculated the annual mean temperature (°C) and annual precipitation (mm) for each year and site. I estimated temperature differences from elevation differences between each of the forest plots and its nearest meteorological station using 0.55°C/100 m in elevation as the mean lapse rate; this relationship between temperature and elevation is commonly used by Japanese forestry researchers (e.g., Oshima et al. 1961, Fukushima et al. 2009). I regressed annual mean temperature and annual precipitation against year for each plot using an ordinary least squares approach in R (R Development Core Team 2019). The resulting regression coefficients indicated the average temperature change per year in the targeted decade, and hence average change rate in annual mean temperature and annual precipitation in each plot. The average rates of change in annual mean temperature (°C) and annual precipitation (mm) were 0.004 ± 0.047 (–0.105 to 0.063) and 5.714 ± 32.795 (–37.082 to 77.791), respectively, in deciduous broad-leaved forests, and 0.003 ± 0.043 (–0.040 to 0.059) and 20.860 ± 18.737 (–0.176 to 47.261), respectively, in evergreen broad-leaved forests [mean ± SD (range)].

### Modeling patterns of existing and novel indices describing community changes across time

I focused mainly on the following indices: *d*_*t.BC*_, *d*_*MR*_, *d*_*M*_/*d*_*R*_, *d*_*E*_/*d*_*S*_, and *v*_*s*_. Formulas and ecological interpretation are summarized in Table 1. First, I analyzed the relationships between these indices and the average rates of change in annual mean temperature and annual precipitation using both simple and polynomial regressions. Then I compared the Akaike’s information criterion (AIC) values (Akaike 1976) among models, including the null model, and selected the best model for each. I did not use multiple regressions for further analyses because the number of data points available was insufficient. All statistical analyses were conducted in R. R scripts to calculate the new indices are provided in Supplementary file 1.

**Table 1.**
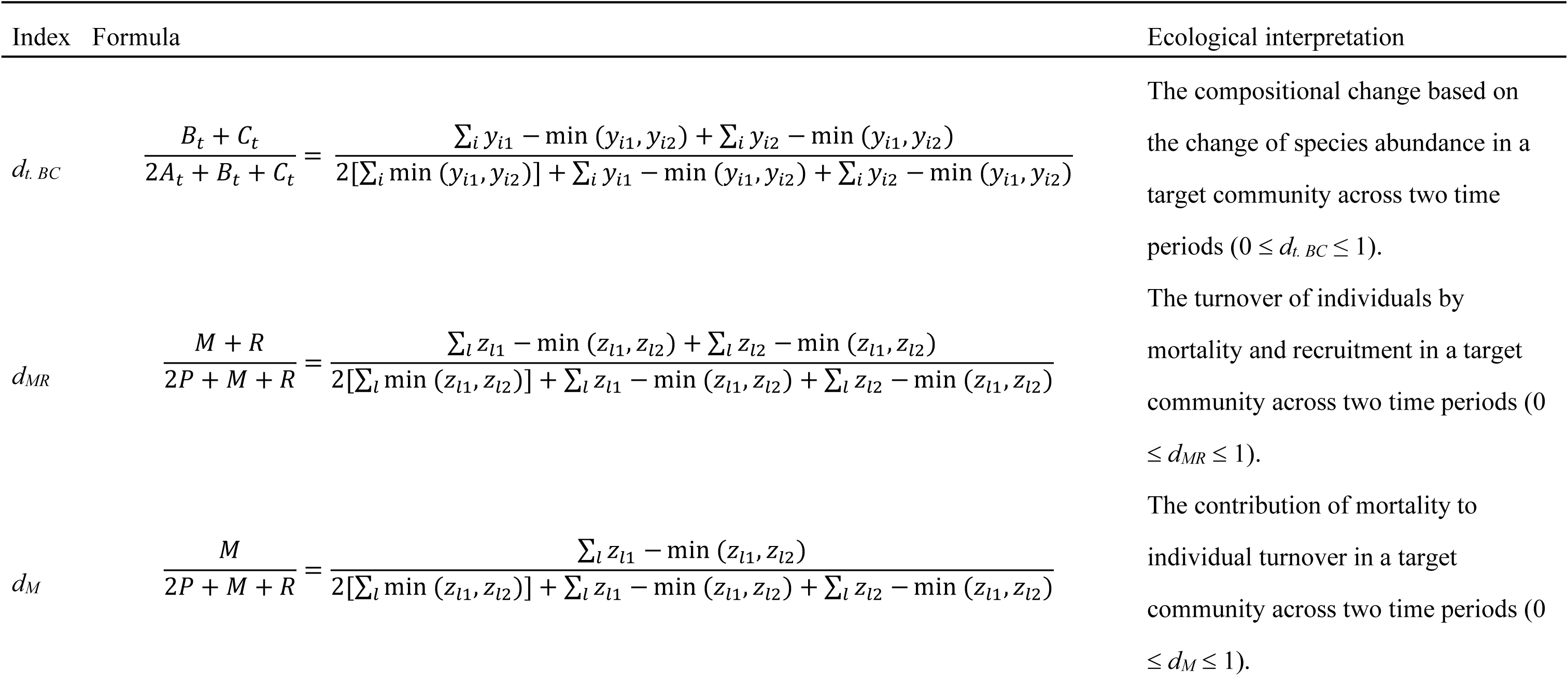
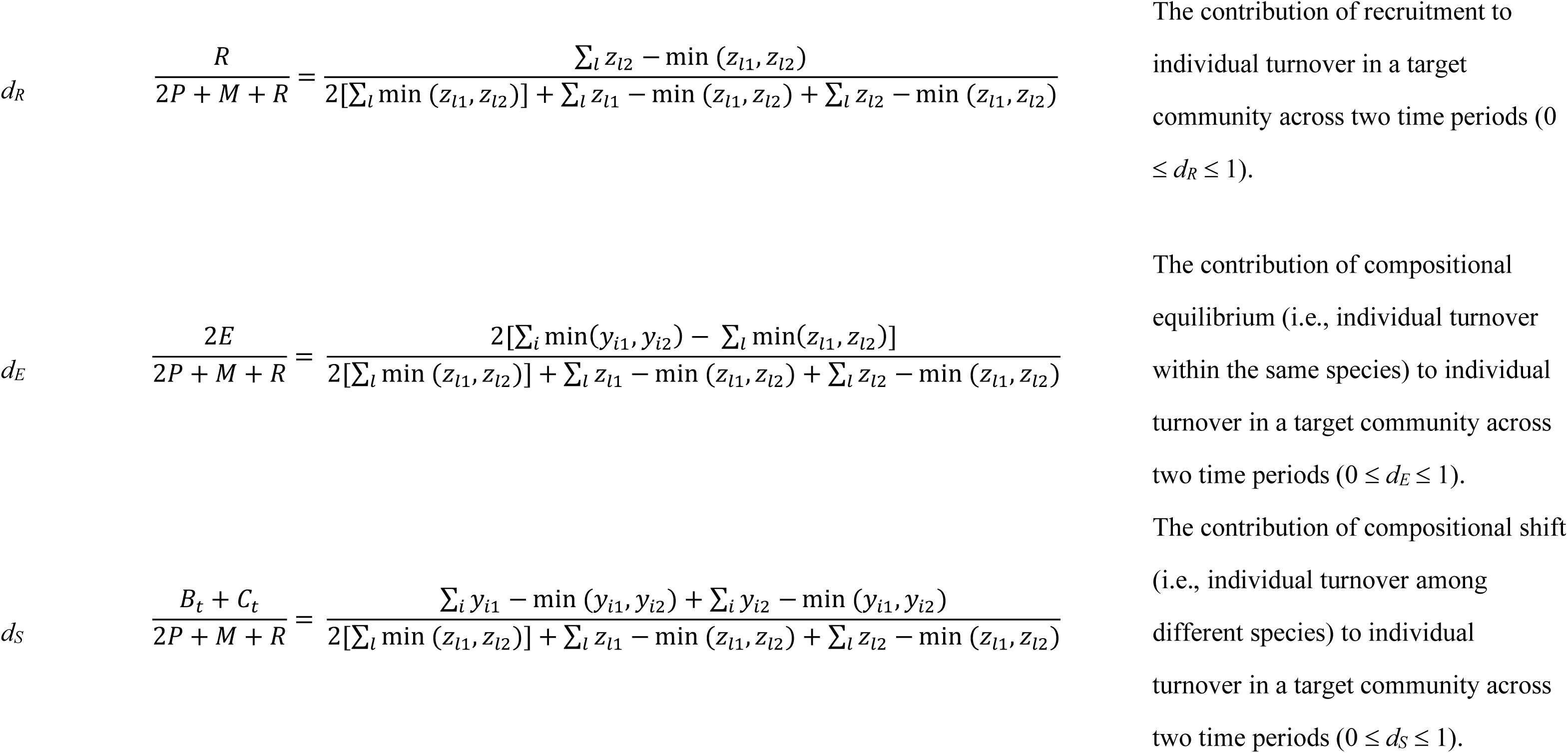
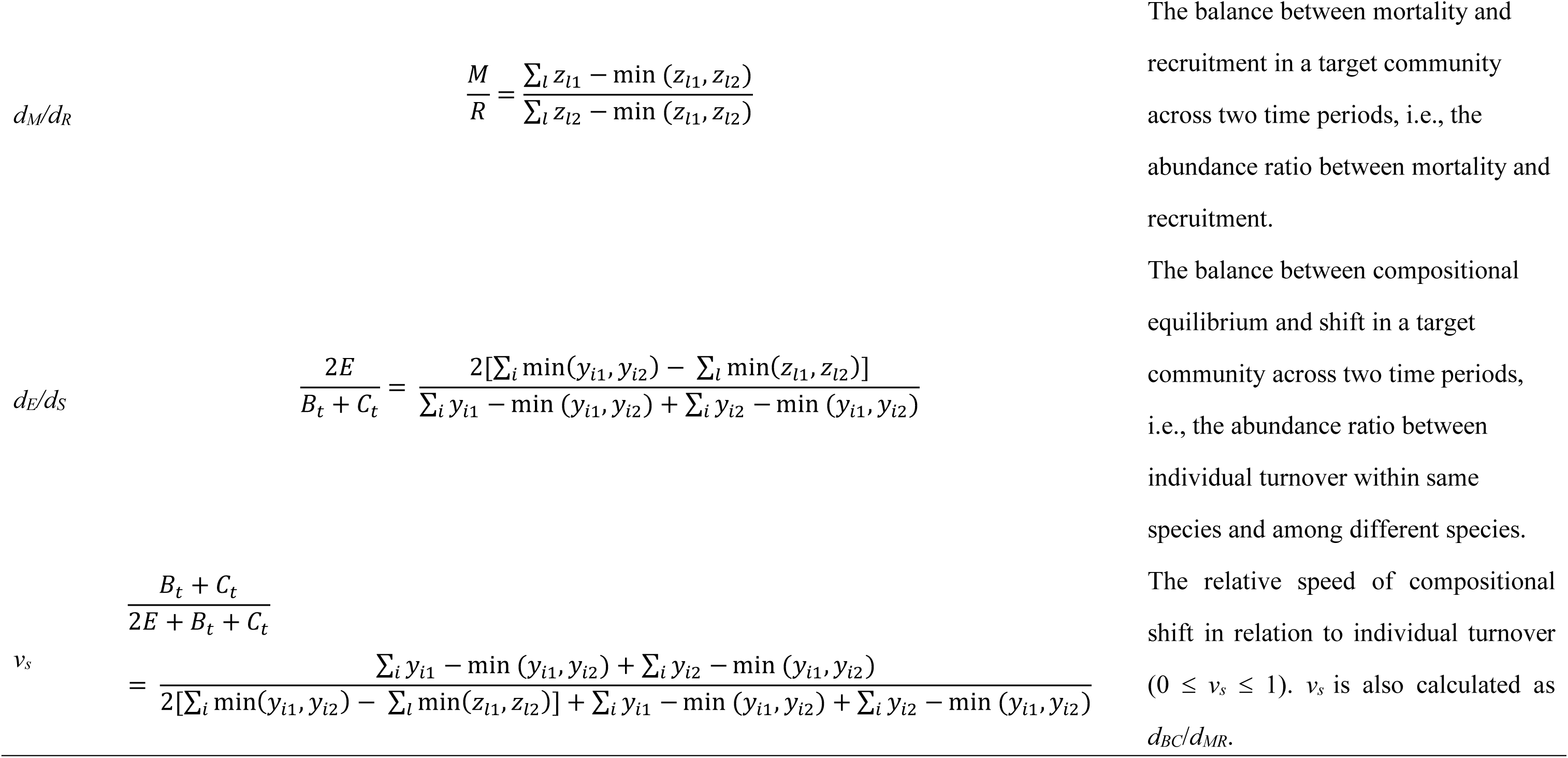
Formulations and ecological interpretation of Bray–Curtis dissimilarity (top: Bray and Curtis 1957) and the novel indices proposed in this study. *y*_*i1*_ is the abundance of species *i* at time T1 at a site, and *y*_*i2*_ is the abundance of species *i* at time T2 at the site. *z*_*l1*_ is individual *l* at time T1, and *z*_*l2*_ is individual *l* at time T2. Both *z*_*l1*_ and *z*_*l2*_ take only *1* or 0, i.e., presence or absence.

## Results

I detected no trends in *d*_*t.BC*_ and *d*_*MR*_ in relation to average rate of change in annual mean temperature in either deciduous or evergreen broad-leaved forests (Fig. 3a, b, c, d, Table S3). In analyses of *d*_*M*_/*d*_*R*_, the polynomial model was chosen by the AIC comparison as the best model for deciduous broad-leaved forests; the simple model was the best for evergreen broad-leaved forests. Values of *d*_*M*_/*d*_*R*_ tended to be greater at sites that warmed the most in the targeted decades. This was true for both forest types. Thus, the relative contribution of mortality to individual based beta-diversity increased with warming rate over time (Fig. 3e, f, Table S3). The *d*_*E*_/*d*_*S*_ index did not show any statistical trends over time for deciduous forests (Fig. 3g, Table S3); the simple regression model was chosen by AIC comparison as the best for evergreen forests (Fig. 3h, Table S3). In evergreen forests, *d*_*E*_/*d*_*S*_ increased most over a decade in warming sites. Thus, the relative contribution of individual turnover without compositional shift in relation to that with compositional shift increased over time (Fig. 3h, Table S3). The polynomial model of *v*_*s*_ was selected as the best for the deciduous forest type: *v*_*s*_ increased with larger temperature changes (Fig. 3i, Table S3). The simple model was selected for the evergreen forest type: *v*_*s*_ decreased linearly with increasing warming over time (Fig. 3j, Table S3). The correlations of all calculated indices are provided in Table S1.

**Figure 3.**
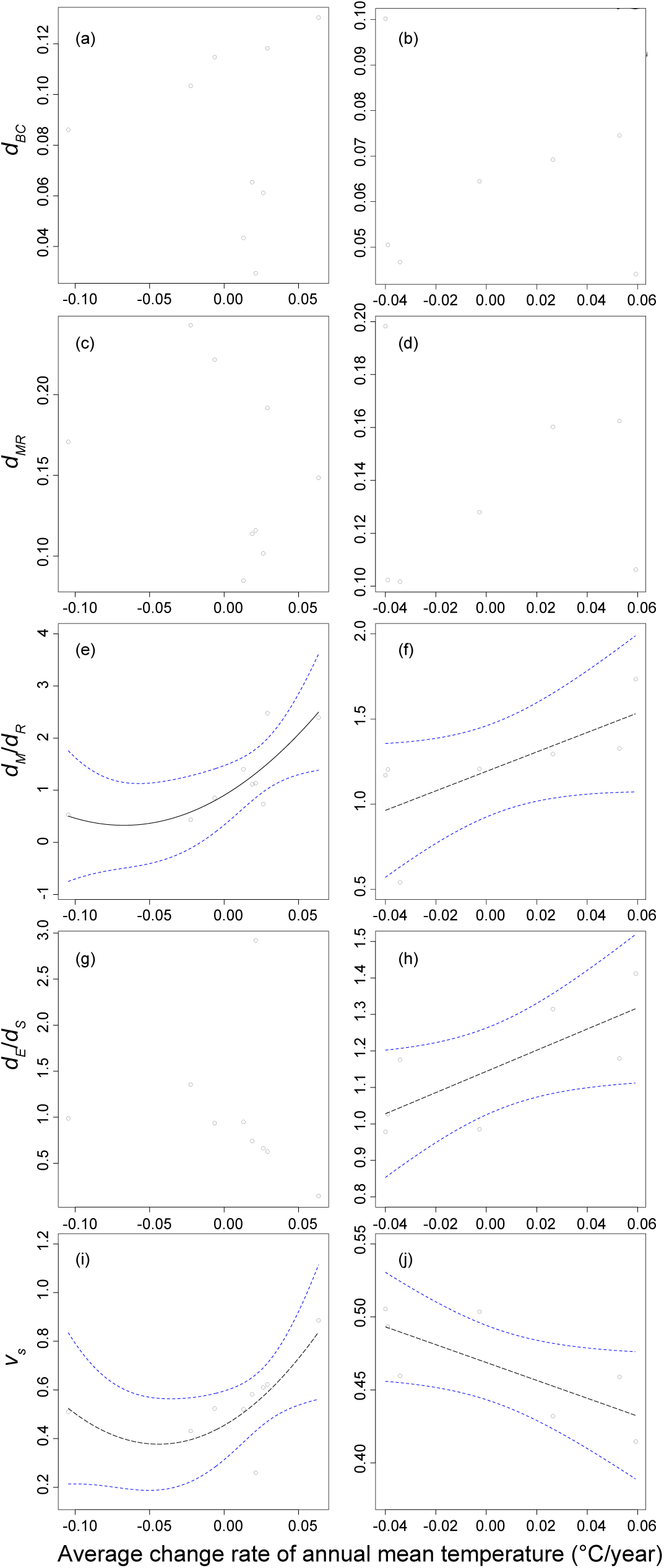
Trends in tree community indices across a gradient of temperature change rates over a decade. The left column of panels refers to trends in deciduous broad-leaved forests; the right column of panels refers to trends in evergreen broad-leaved forests. Panels (a) and (b) show the trends in typical abundance-based temporal beta-diversity indices (Bray–Curtis dissimilarity: *d*_*BC*_). Panels (c) and (d) show the trends in individual-based temporal beta-diversity indices (*d*_*MR*_). Panels (e) and (f) show trends in the *d*_*M*_/*d*_*R*_ index, which is an indicator of the relative contribution of mortality and recruitment processes to individual-based temporal beta-diversity (e: *adjusted-R*^*2*^ = 0.523, *P* = 0.046; f: *adjusted-R*^*2*^ = 0.391, *P* = 0.079). Panels (g) and (h) show trends in the *d*_*E*_/*d*_*S*_ index, which is an indicator of the relative contribution of equilibrium and shift processes to individual-based temporal beta-diversity (h: *adjusted-R*^*2*^ = 0.472, *P* = 0.053). Panels (i) and (j) show trends in the index *v*_*s*_, which is an indicator of the relative speed of compositional shift in relation to individual turnover (i: *adjusted-R*^*2*^ = 0.413, *P* = 0.085; j: *adjusted-R*^*2*^ = 0.462, *P* = 0.056). The solid black lines indicate statistically significant (*P* < 0.05) trends. The dashed black line indicates a marginally significant (0.05 < *P* < 0.1) trend. The blue dashed lines enclose the 95% confidence interval envelopes. Detailed results of statistical analyses are presented in Table S3.

The average rate of change in annual precipitation did not affect trends over time for any of the calculated indices (Table S3).

## Discussion

The effects of anthropogenic climate change have been intensively studied in many disciplines, e.g., analyses of species interactions, range shifts, phenology, and ecosystem functioning (Thomas et al. 2004, Kordas et al. 2011, Valtonen et al. 2014). However, studies of temporal changes in diversity and composition in local communities have often produced mixed and/or contradictory results among sites and researchers (Vellend et al. 2013, Dornelas et al. 2014, Newbold et al. 2015, reviewed in Hisano et al. 2018). Quantitative analyses of biological community change over time are required to expand our understanding of global climate change and its relationship to biodiversity change (Dornelas et al. 2014). The novel concepts and procedures that I have developed quantitatively evaluated the relative speed of compositional shift and the relative contribution of mortality and recruitment processes in tree communities along a gradient in temperature change rates across a decade. These patterns had previously been difficult to detect using existing abundance-based dissimilarity indices (Fig. 3a, b). Compositional shifts in tree communities are generally recognized as long-term phenomena in the absence of disturbance. Therefore, 10 years is a relatively short-time period in which to analyze compositional changes in tree communities comprising species with life spans extending from decades to centuries (Loehle 1988). I found that community responses to local climate change trends differed between forest types, but nevertheless, there were real changes in community composition over only a decade. These changes were associated with change in annual mean temperatures, but not with changes in annual precipitation across the Japanese archipelago.

One advantage of the method I have developed is that it dissects individual turnover into mortality and recruitment components. For example, I detected no clear patterns in the relationship between individual turnover (*d*_*MR*_), which is the obligate speed of temporal change in communities, and temperature change rate (Fig. 3c, d). However, by splitting the index into two components, I was able to identify relative contributions to individual turnover (*d*_*M*_*/d*_*R*_). When this was done, clear patterns emerged showing that an increase in the temperature change rate facilitated the relative contribution of mortality components (Fig. 3e, f). Furthermore, by assessing the trend of each component separately, I was able to confirm that decreases in recruitment components with increases in the temperature change rate drove the pattern in deciduous forests, but I was not able to confirm this in evergreen forests (Table S4).

The relative contribution of the index *d*_*E*_*/d*_*S*_ to individual-based temporal beta-diversity (*d*_*MR*_) increased positively in evergreen forests as the warming rate increased, although no trends in the individual components (*d*_*E*_, *d*_*S*_) were discerned. Thus, the balance between the components changed in a consistent manner with temperature change, although no patterns of variation in the individual components were found. There were no clear trends in *d*_*E*_*/d*_*S*_ in relation to temperature change rates in deciduous forests. However, *d*_*E*_ was affected by the temperature change rate in these forests: *d*_*E*_ decreased with increased warming rates; thus, there was a clear contribution of compositional equilibrium decrease along these gradients (Table S4).

I used a novel index to evaluate compositional changes, i.e., the relative speed of the compositional shift in communities (*v*_*s*_), which clearly changed along the gradient of temperature change rates (Fig. 3i, j, Table S3). Furthermore, the responses were different between deciduous and evergreen broad-leaved forests. Specifically, the relative speed in evergreen forests was smaller at sites with greater temperature warming. Furthermore, the range size of the values (*v*_*s*_) in deciduous forests was about 7-fold greater than the range size in evergreen forests (Table S5). Thus, deciduous forests are more affected by differences in temperature change rate. I did not detect statistically significant trends along a gradient of precipitation change rate, although climate change induced drought is a major factor in tree mortality (Allen et al. 2010, Saiki et al. 2017). These contrasting results may be due to the humid climate of the Japanese archipelago, but assessing this relationship requires the inclusion of many other candidate explanatory variables (e.g., continuous dry days) which were not available. To capture the effects of global climate change accurately, a broad multi-site analysis across the globe using similar methods and other relevant environmental variables will be required.

Predicting community persistence from snapshot data on community structure is a complicated issue for community ecologists. For example, Ulrich et al. (2018) argued that it may be possible to evaluate species persistence and their relative proportions from the patterns of species abundance distribution. If this were the case, the relative speed of compositional shift should provide a comparable estimate of community persistence from both individual and species-based perspectives. Specifically, using some time series datasets containing substantial annual individual information, it should be technically possible to calculate the retention time for each community, i.e., the lifespan durations of all individuals replaced in a focal community. These calculations could be cross-compared with snapshot community structure data (e.g., species abundance patterns). In the present study, I used data for only two time periods, so this cross-comparison was not possible. Nevertheless, clarification of the relationship between relative abundance and temporal changes in individual and community composition should become possible in the near future. Underlying mechanisms explaining these relationships may then be discernible.

Compilation of long-term temporal datasets for large macroscopic organisms (e.g., trees and mammals) requires considerable human effort and cost for monitoring and research surveys because large organisms often have long life spans. Nevertheless, the new indices I developed clearly highlighted the importance of monitoring datasets assembled using traditional detailed surveys (e.g., individual tracking in forest plots) in detecting the temporal trends over short time periods. Therefore, I re-emphasize here the significance of classical monitoring surveys to collect biodiversity information. I also developed an approach that introduced the concept of individual-based beta-diversity within the context of temporal changes in community composition. Theoretically, however, the procedure could be expanded to a spatial context for use in studies of animal behavioral ranges and movements, including long-distance migration, using monitoring and biologging datasets (Viana et al. 2016, Börger et al. 2020) that include multiple time series datasets collected from the same place within a shorter time frame than the life spans of the targeted organisms (e.g., bird ringing and classical mark–recapture procedures). Specifically, the approach could detect detailed spatial turnover for individuals within same species in communities.

Many ecology theories have been developed from the concept of species. The original concept of beta-diversity was proposed to reflect species turnover (Whittaker 1960). By contrast, much recent research has pointed out the problems of species-based studies and the importance of considering individuals in community ecology (Dupont et al. 2011, Nakazawa 2017, Tatsumi et al. 2019). Species are the consequences of ecological/evolutionary processes among individuals; to be accurate, the species themselves do not drive these processes, i.e., individuals do by competing with other individuals, and this does not involve the species directly. Although individuals were considered in the unified neutral theory (Hubbel 2001) and subsequent related studies (Hubert et al. 2015, Vanoverbeke et al. 2016, Tatsumi et al. 2019), the focus was restricted to the patterns of species richness, abundance, and related diversity indices. The effects of individual births and deaths on biodiversity patterns have not yet been considered sufficiently. Procedures for incorporating individual identity information into biodiversity evaluation, i.e., individual-based diversity indices, are required to facilitate further conceptual advances in ecology. New ecological concepts will emerge from combined considerations of species abundance and information relevant to individual identity in the future.

## Conclusions

Previous studies have often detected temporal changes in local communities, likely as a result of human activities, but not at all times and places. In this study, I clearly showed the responses of tree communities to local and short-term climate change trends using datasets collected according to novel index concepts. Notably, the relative speed of compositional shift in tree communities varied along a gradient of temperature change rates over a single decade. This community level response resulted from individual-level mortality and recruitment processes. Therefore, accumulating more detailed long-term information at global scales, specifically individual identity information at the local community level, is required for understanding both actual biodiversity responses to global climate change and background processes. Mortality and recruitment rates are generally difficult to measure in empirical studies compared to loss and gain information based only on snapshot data, especially in the case of plants or vertebrates. The individual-based approach may not be tractable for all taxonomic groups. However, information on individual organisms will help to bridge the disciplines of macroecology and local studies focusing on within-individual phenomena, for example, physiological ecology. The calculation of individual-based diversity indices from individual identity monitoring data will support efforts to quantitatively evaluate the impacts of climate change on future biodiversity. Increasing availability of monitoring data with individual identification combined with individual-based diversity indices such as presented here can thus be key in quantitatively evaluating the impacts of climate change on biodiversity over time.

## Data accessibility statement

All data were obtained from open sources. Forest monitoring data were obtained from the Biodiversity Center of Japan (http://www.biodic.go.jp/moni1000/index.html, in Japanese: accessed on November 11, 2019) published in part by Ishihara et al. (2011). Climate data were obtained from the Japanese Automated Meteorological Data Acquisition System (AMeDAS) database (http://www.jma.go.jp/jma/indexe.html, Japan Meteorological Agency accessed December 12, 2019). Taxonomic information was obtained from the Ylist (Yonekura and Kajita 2003: http://ylist.info/ylist_simple_search.html, in Japanese: accessed December 8, 2019).

## Acknowledgments

I thank Drs. Anu Valtonen and Shunsuke Matsuoka who made insightful comments on an early version of the manuscript and Dr. Takaya Iwasaki who advised on the data visualizations. I also thank my friend for inspiring me to reconsider the importance and the preciousness of individual life in both society and ecology.

## Appendices

**Table S1.** Correlations among geographical and environmental variables and their respective calculated indices.

**Table S2.** Geographical and climate attributes of each plot and the nearest metrological station. See Fig. 2 for plot locations.

**Table S3.** Statistical analyses of the relationships between the indices *d*_*BC*_, *d*_*MR*_, *d*_*M*_/*d*_*R*_, *d*_*E*_/*d*_*S*_, and *v*_*s*_ and the change rate of annual mean temperature and annual precipitation across a decade for two different forest types. The four indices are defined in the legend of Fig. 3. The analyses were performed using three models (null, simple regression, polynomial regression). *adj. R*^*2*^, adjusted coefficient of determination. AIC, Akaike’s information criterion. Fig. 3 presents plots of the trends analyzed in part A of this table.

**Table S4.** Results of simple linear regression analyses of other calculated indices, which were not included in the main analyses, on the change rates of annual mean temperature and annual precipitation.

**Table S5.** The values (mean, standard deviation, and range) of calculated indices for deciduous broad-leaved forest and evergreen broad-leaved forest in the present study.

**Table S6.** Results of simple linear regression analyses of the calculated indices using latitude and the means and SDs of annual mean temperature and annual precipitation

**Table S7.** Indices calculated in this study

**Figure S1.** Annual mean temperature and annual precipitation in each year (2003–2018) at each site. The red circles and lines indicate the annual mean temperature; the blue bars indicate annual precipitation. The dashed line indicates the average annual mean temperature (1981–2010). The dot-dash line indicates the average of annual precipitation (1981–2010).

**Supplementary text 1.** Explanation of individual-based temporal beta-diversity in the Ružička dissimilarity index version.

**Supplementary file 1.** R software script of individual-based temporal beta-diversity calculations using sample data

