## Supplementary figures and images for "Degrees of compositional shift in tree communities vary along a gradient of temperature change rates over one decade: Application of an individual-based temporal beta-diversity concept"

### Figure S1

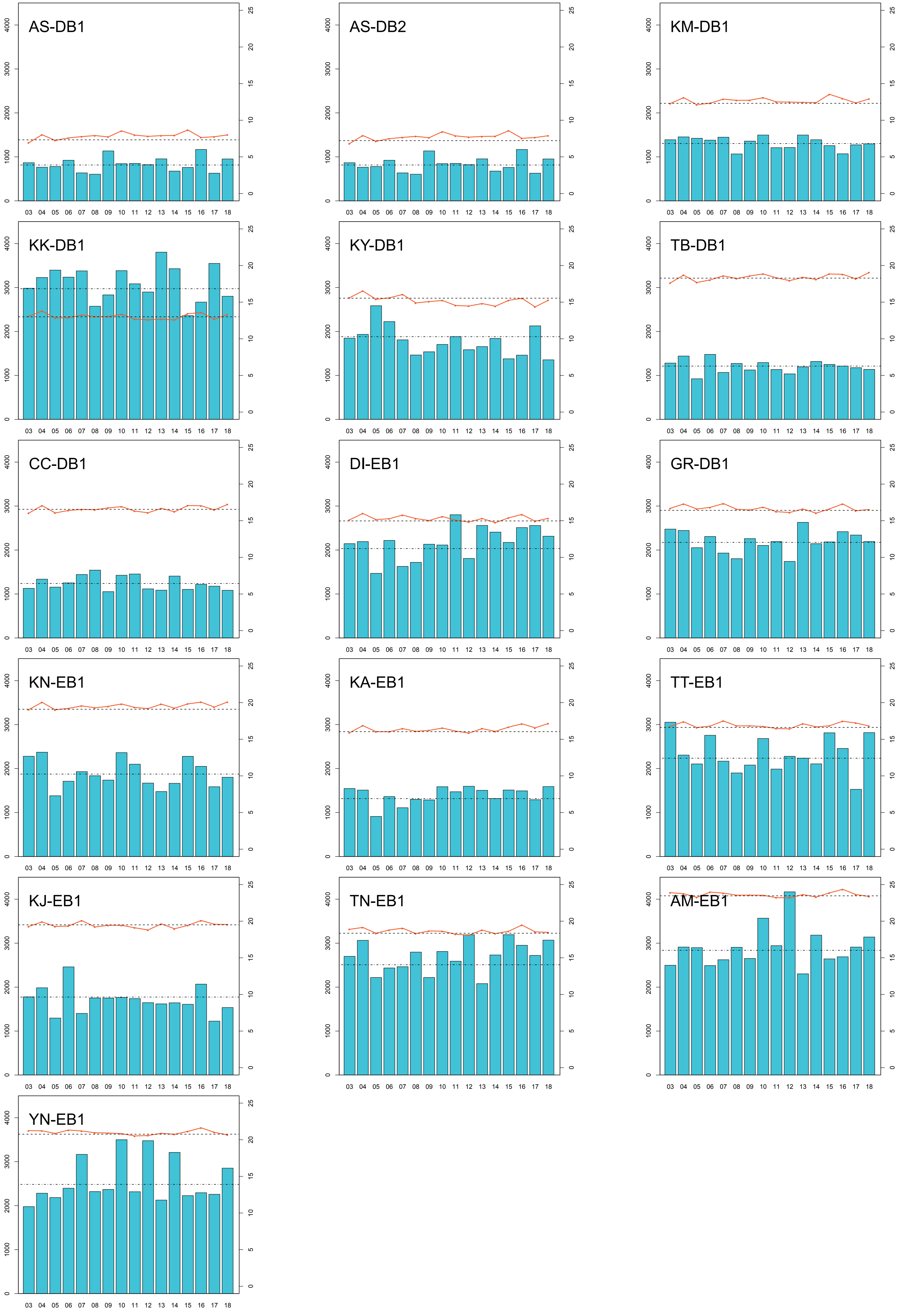
