## Supplementary file 1 for "Degrees of compositional shift in tree communities vary along a gradient of temperature change rates over one decade: Application of an individual-based temporal beta-diversity concept": function_example_script.R

```
#convertion from individual information to demographic information
convert_indiv_demo <- function(x){
  x <- data.frame(x)
  
  t1 <- table(x[,2][x[,3]==1])
  t2 <- table(x[,2][x[,4]==1])
  m <- table(x[,2][x[,3]==1&x[,4]==0])
  r <- table(x[,2][x[,3]==0&x[,4]==1])
  
  t1_d <- data.frame(t1)
  names(t1_d) <- c("species","t1")
  t2_d <- data.frame(t2)
  names(t2_d) <- c("species","t2")
  m_d <- data.frame(m)
  names(m_d) <- c("species","m")
  r_d <- data.frame(r)
  names(r_d) <- c("species","r")
  
  merge2 <- function(dfs, ...) # original merge2 script from https://gist.githubusercontent.com/yuu-ito/7094938/raw/336a780aa0d91c3cbdbb5e6c747b244a2c83ed40/002_merge2.R
  {
    base <- dfs[1]
    lapply(dfs[-1], function(i) base <<- merge(base, i, ...)) 
    return(base)
  }
  
  conv_data <-   merge2(list(t1_d,t2_d,m_d,r_d),by="species",all=T)
  conv_data[is.na(conv_data)] <- 0
  conv_data[2:5] <- apply(conv_data[2:5],2,as.numeric)
  return(conv_data)
}


#calculation of individual-based temporal beta diversity 
beta.indiv<-function (x,index.family) {
  
t1 <- as.matrix(x[2])
t2 <- as.matrix(x[3])
m <- as.matrix(x[4])
r <- as.matrix(x[5])
  
T1 <- sum(t1)
T2 <- sum(t2)
M <- sum(m)
R <- sum(r)

A <- sum(pmin(t1,t2))
E <- sum(pmin(m,r))
B <- M - E
#B <- sum(t1)-sum(pmin(t1,t2))
C <- R - E
#C <- sum(t2)-sum(pmin(t1,t2))
P <- A - E

index.family <- match.arg(index.family, c("Bray_Curtis", "Ruzicka"))

switch(index.family, "Bray_Curtis" = {

#Bray-Curtis version

      BC <- (B+C)/(2*A+B+C) 
      BC_loss <- B/(2*A+B+C) 
      BC_gain <- C/(2*A+B+C) 
      MR <- (M+R)/(2*P+M+R)
      MR_mortality <- M/(2*P+M+R) 
      MR_recruitment <- R/(2*P+M+R) 
      MR_equilibrium <- (2*E)/(2*P+M+R)
      MR_shift <- (B+C)/(2*P+M+R)
      vs <- (B+C)/(2*E+B+C) 

      results <- c(BC,BC_loss,BC_gain,MR,MR_mortality,MR_recruitment,MR_equilibrium,MR_shift,vs)
      names(results) <- c("BC","BC_loss","BC_gain","MR","MR_mortality","MR_recruitment","MR_equilibrium","MR_shift","vs")
      
}, "Ruzicka" = {
      
#Ruzicka version     
      
      BC <- (B+C)/(A+B+C) 
      BC_loss <- B/(A+B+C)
      BC_gain <- C/(A+B+C) 
      MR <- (M+R)/(P+M+R)
      MR_mortality <- M/(P+M+R)
      MR_recruitment <- R/(P+M+R) 
      MR_equilibrium <- (2*E)/(P+M+R) 
      MR_shift <- (B+C)/(P+M+R) 
      vs <- (B+C)/(E+B+C)
      
      results <- c(BC,BC_loss,BC_gain,MR,MR_mortality,MR_recruitment,MR_equilibrium,MR_shift,vs)
      names(results) <- c("BC","BC_loss","BC_gain","MR","MR_mortality","MR_recruitment","MR_equilibrium","MR_shift","vs")
      
})
      
  return(results)

}

###example#######

sample_data <- read.csv("sample_data.csv")

test <- convert_indiv_demo(sample_data)

beta.indiv(test,"Bray_Curtis")

beta.indiv(test,"Ruzicka")
```
